# Phylogenetic and evolutionary analysis reveals the recent dominance of ciprofloxacin resistant *S. sonnei* and local persistence of *S. flexneri* clones in India

**DOI:** 10.1101/2020.06.05.137372

**Authors:** Dhiviya Prabaa Muthuirulandi Sethuvel, Ankur Mutreja, Agila Kumari Pragasam, Karthick Vasudevan, Dhivya Murugan, Shalini Anandan, Joy Sarojini Michael, Kamini Walia, Balaji Veeraraghavan

**Author notes:** Corresponding author, Dr. Balaji Veeraraghavan, Professor, Dept. of Clinical Microbiology, Christian Medical College, Vellore 632004, Tamil Nadu, India.

## Abstract

*Shigella* is the second leading cause of bacterial diarrhea worldwide. Recently *S. sonnei* seems to be replacing *S. flexneri* in low and middle-income countries undergoing economic development. Despite this, studies focusing on these species at genomic level remain largely unexplored. Here we compared the genome sequences of *S. flexneri* and *S. sonnei* isolated from India with the publically available genomes of global strains. Our analysis provide evidence for the long term persistence of all PGs of *S. flexneri* and the recent dominance of ciprofloxacin-resistant *S. sonnei* lineage in India. Within *S. flexneri* PGs, majority of the study isolates belonged to PG3 within the predominance of serotype 2. For *S. sonnei*, the current pandemic involves globally distributed MDR clones that belong to Central Asia lineage III. The presence of such epidemiologically-dominant lineages in association with stable AMR determinants results in the successful survival in the community.

## Introduction

*Shigella* is ranked as the second leading cause of bacterial diarrhea worldwide and the third leading cause of deaths in children less than 5 years (Kotloff, 2018). This has been categorized as a priority pathogen among enteric bacteria on the Global Antimicrobial Resistance Surveillance System (GLASS) by the World Health Organization (WHO) (WHO, GLASS report, 2017-2018). Among the four species of *Shigella*, the most common cause of endemic shigellosis, and the most frequently isolated species in India is *S. flexneri*. However, *S. sonnei* seems to be replacing *S. flexneri* in low and middle-income countries undergoing economic development. *S. dysenteriae* and *S. boydii* are now relatively uncommon (Thompson et al., 2015). The reasons behind this changing species level epidemiology remain unclear. Nevertheless, the increasing awareness of the disease burden and the emerging threats posed by drug resistant *Shigella* have resulted in continued interest in the development of *Shigella* vaccines, for which several candidates are currently undergoing testing in clinical trials.

Antimicrobial resistance in *S. flexneri* and *S. sonnei* is a growing international concern, specifically with the international dominance of the multi-drug resistant (MDR) lineage of *S. sonnei* (The HC et al., 2016). Earlier phylogenetic analysis predicted that South Asia would be the hub for the international spread of ciprofloxacin-resistant *S. sonnei* (The HC et al., 2016). However, only a limited number of *Shigella* isolates from India have been included in previous studies. Therefore, understanding the expansion of this lineage and the phylogenetic relationship with the isolates from within and outside Asia, including India is critical.

Genomic studies focusing on *S. flexneri* and *S. sonnei* in India remain largely unexplored. Understanding the genome sequences of antimicrobial-resistant pathogens can enhance our knowledge of the molecular identity of resistance traits and their mechanism of dissemination within the microbial population. Here we compared the genome sequences of *S. flexneri* and *S. sonnei* isolated from India with the publically available genomes of global strains, to understand the introduction and expansion of drug-resistant strains in India. Bayesian phylogenetic analysis was performed, in particular for *S. sonnei* isolates from India to demonstrate the evolution of ciprofloxacin resistant *S. sonnei* clones in India. In addition, virulence, resistance and plasmid profiles of the isolates were analyzed and correlated with the previously defined *S. flexneri* phylogenetic groups and *S. sonnei* lineages.

## Results

The whole-genome phylogenetic tree was constructed using 106 *S. flexneri* and 82 *S. sonnei* genomes sequenced in this study and 60 *S. flexneri* and 362 *S. sonnei* genomes from previous studies. All *S. flexneri* and *S. sonnei* sequences were mapped to the reference sequences (Accession numbers: NC_007384 and NC_004337.2).

### Phylogenetic analysis of *S. flexneri*

The phylogenetic comparison of Indian *S. flexneri* isolates against isolates available from other South Asian countries (Pakistan and Bangladesh) showed that Indian isolates were distributed across all the previously defined phylogenetic groups (PG1, 2, 3, 5, 6, 7) with the exception of PG4 (**Fig 1**). The majority of Indian isolates clustered within PG3 (70%), followed by PG1 (22%) and PG2 (7%). Each PG contained multiple serotypes. However, the common serotypes found within PG2, PG3 and PG5 was *S. flexneri* serotype 3a, 2a and 5a respectively. Variation in the distribution of virulence determinants among the PGs was also evident. Notably, the SHI-1 pathogenicity island (PAI), which is known to carry *pic, sigA* and ShET-1 – *set*1A/B genes was exclusively seen in PG3 isolates with exception of a single isolate of PG1. SHI-1 was present in various serotypes within PG3 but was predominantly seen in serotype 2a. Additionally, SHI-2 PAI known to carry *iuc*ABCD and *iutA* genes was found in the isolates of all PGs. Similarly, *sat*, a serine protease autotransporter was observed in all PGs. The *sen* gene encoding enterotoxin ShET-2 was only seen in the PG2, and PG7 isolates. Enterobactin genes (*ent*BCEFS and *fep*ABCDG) that encode iron siderophore were present across PG3, PG4, PG5, PG7, and were also in a few isolates of PG1. Notably, *fimE* gene was absent in all PG3 isolates except one.

**Fig 1:**
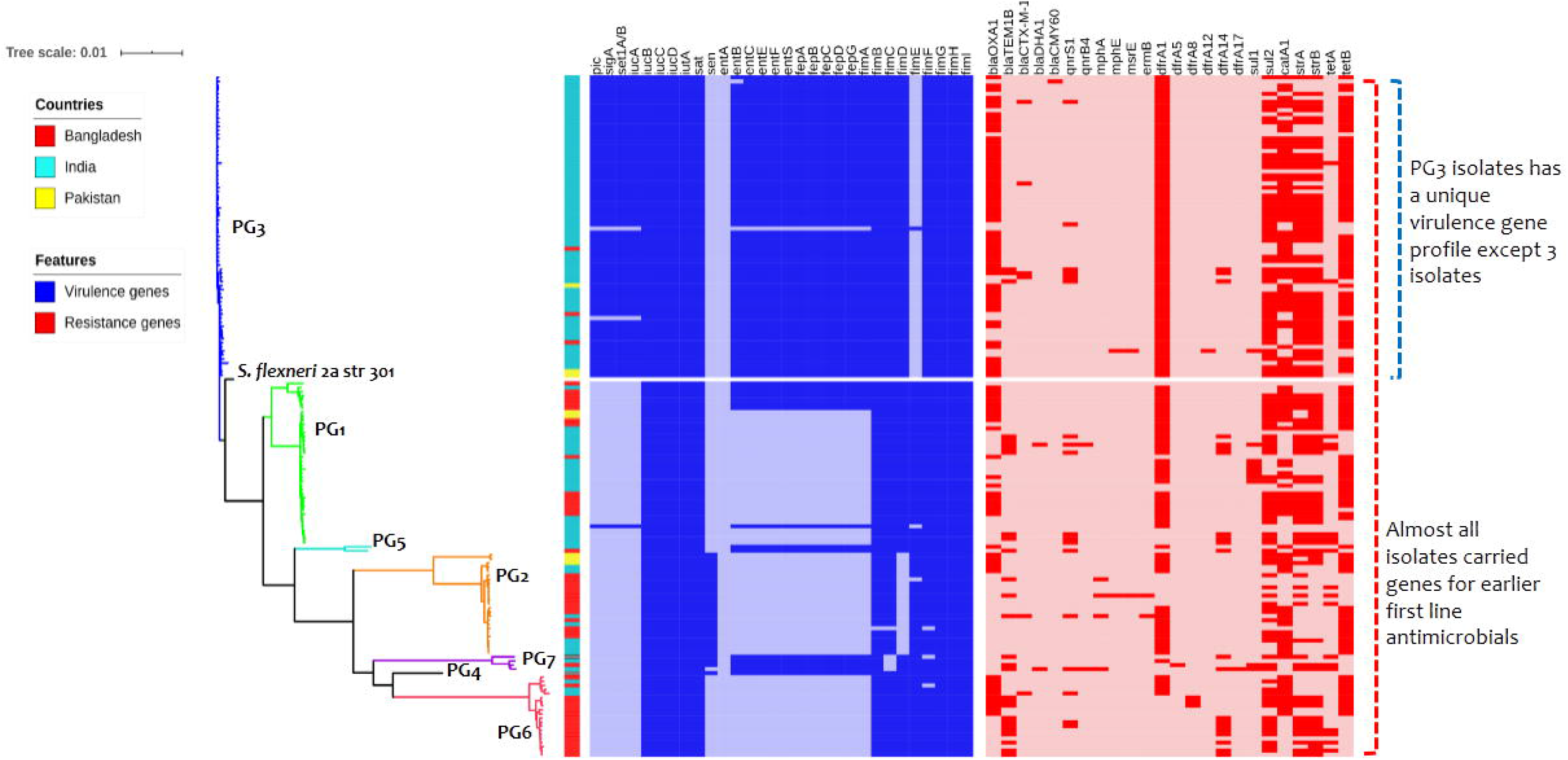
A Maximum-likelihood phylogenetic tree of 166 *S. flexneri* isolates mapped against the *S. flexneri* 2a str 301 reference genome. The clade colors represent to the previously described phylogenetic groups (PGs). Virulence genes are highlighted in blue and resistance genes are highlighted in red. The scale bar indicates the number of substitutions per site.

Screening of known AMR genes revealed highly variable AMR gene distribution both within and across PGs (**Fig 1**). Genes encoding resistance to earlier first-line antimicrobials includes, streptomycin (*str*A/B), Trimethoprim/sulfamethoxazole (*dfrA1, sul)*, chloramphenicol (*cat*A1) and tetracycline (*tet*A/B) were distributed across all the PGs. Among beta-lactamases, the *bla*_OXA-1_ gene was present in all PGs with the exception of PG4 and PG7. Similarly, *bla*_TEM-1B_ was present across all PGs with the exception of PG4, whereas *bla*_CTX-M-15_ was only seen in PG2, PG3, and PG6. The AmpC beta-lactamase genes, *bla*_DHA_ was identified in two isolates, one each from PG1 and PG7. In addition, the *bla*_CMY-60_ gene was identified in PG3 alone. These AmpC genes were only observed in the Indian isolates. Furthermore, the analysis of fluroquinolone resistance mechanisms showed the presence of both plasmid mediated quinolone resistance (PMQR) genes and mutations within the quinolone resistance determining region (QRDR). The PMQR gene, *qnrS1*, was identified in all PGs with the exception of PG4 whilst *qnrB4* was seen in two isolates, one each from PG1 and PG7. The most frequently observed QRDR mutations were S83L in *gyrA* and S80I in *parC* as a result of replacement of serine by leucine and isoleucine, respectively. Macrolide resistance genes were recognized, as *mphA* in PG2; and PG7, *ermB* and *mphE* in PG2; and *msrE* in PG3.

Additional plasmid analysis showed that, IncFII plasmid was the most predominant across the PGs followed by ColRNAI (**Fig 1**). Among the Inc plasmid types, IncX3 was found in isolates belonging to PG1, PG2 and PG3, whereas IncY/IncL/M/IncI2 were only identified in PG3 isolates. Furthermore, IncFIC was only seen in PG1 whilst IncFIB was seen in all PGs with the exception of PG4.

### Phylogenetic analysis of *S. sonnei*

The phylogenetic analysis of *S. sonnei* revealed that the ciprofloxacin-resistant isolates were grouped within the Central Asia III lineage. The majority of the isolates in the Central Asia III lineage carried genes encoding resistance to streptomycin, trimethoprim/sulfamethoxazole and tetracycline as described earlier. Most of the isolates (69%) within this lineage carried triple mutation (*gyr*A – S83L, D87G/N, *par*C – S80I) in the QRDR regions (**Fig 2**). Double and single mutation was observed in 1% and 19% of the isolates. Only one isolate carried PMQR, *qnr*B gene. The Central Asia Lineage III had a different plasmid profile compared to the isolates of other lineages. Other lineages carried ciprofloxacin susceptible isolates with varying AMR and plasmid profiles. In addition, analysis of virulence determinants among the *S. sonnei* isolates showed a varying profile of virulence genes in comparison to *S. flexneri*. Notably, *S. sonnei* isolates were found to only carry the *sig*A gene within the SHI-1 PAI whilst harbouring multiple virulence genes within the SHI-2 PAI. Further, enterotoxin (*sen*) and enterobactin (*ent*BCHFS and *fep*ABCDG) genes were identified among the isolates.

**Fig 2:**
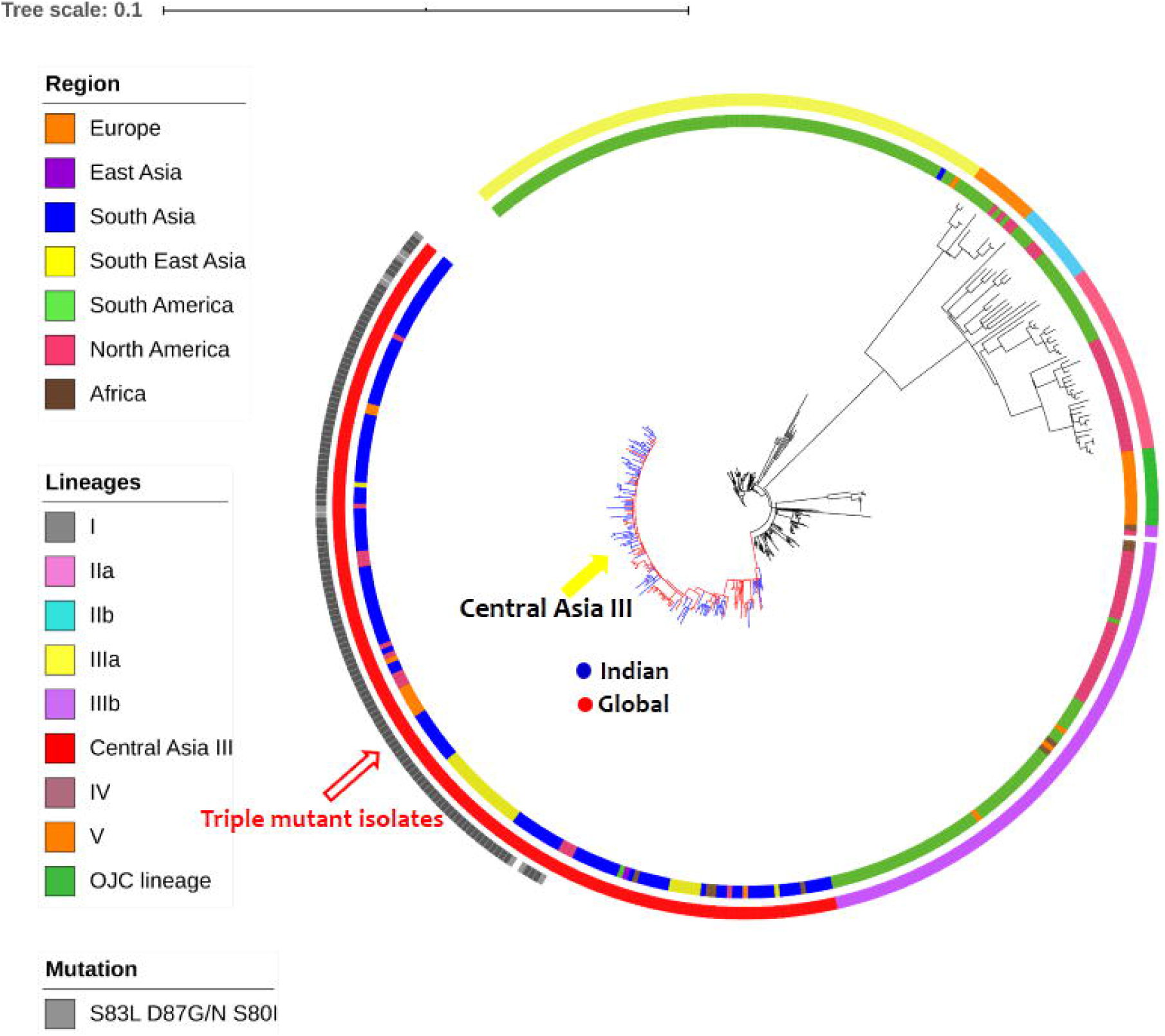
A Maximum-likelihood phylogenetic tree of the Vellore ciprofloxacin resistant *S. sonnei* in a global context. The tree includes 448 *S. sonnei* sequences including reference *S. sonnei* Ss046. The clade color represents the Central Asia III lineage (India – blue, global – red). The inner ring represents the region of the isolates followed by lineages of *S. sonnei* and the outer ring indicates QRDR mutation (triple mutant isolates). The scale bar indicates the number of substitutions per site.

As there was a strong temporal signature for the observed mutations in the global *S. sonnei* population, we explored the temporal structure of the *S. sonnei* isolates that belonged to Central Asia III lineage from India using Bayesian phylogenetic methods. The root to tip analysis revealed a strong correlation (R squared 0.7543) between the time of isolation and distance from root suggesting the temporal clock like evolution in the lineage (**Fig S1**). The median substitution rate of the *S. sonnei* population was estimated to be 2.083×10-3 substitutions per base per year in this study. This time-scaled phylogenetic reconstruction demonstrated the sequential accumulation of QRDR mutations among the *S. sonnei* population. The most recent common ancestor (MRCA) of the ciprofloxacin resistant triple mutant clade in India was estimated to be from the year 2000 (95% HPD: 1984 – 1987). Bayesian hierarchical clustering using core SNPs segregated the ciprofloxacin-resistant isolates that had a triple mutation in QRDR into a separate clade (**Fig 3**). The mutations, *gyr*A-S83L was estimated to occur in ~1996 (95% HPD: 1995–2001.1), *gyr*A-D87Y in ~1996 (95% HPD: 1996–1999.8) and triple mutations S83L, D87Y/G and S80I in ~2003 (95% HPD: 2003–2006.8). This infers that the ciprofloxacin-resistant population expanded rapidly after 2004 and has since sustained.

**Fig 3:**
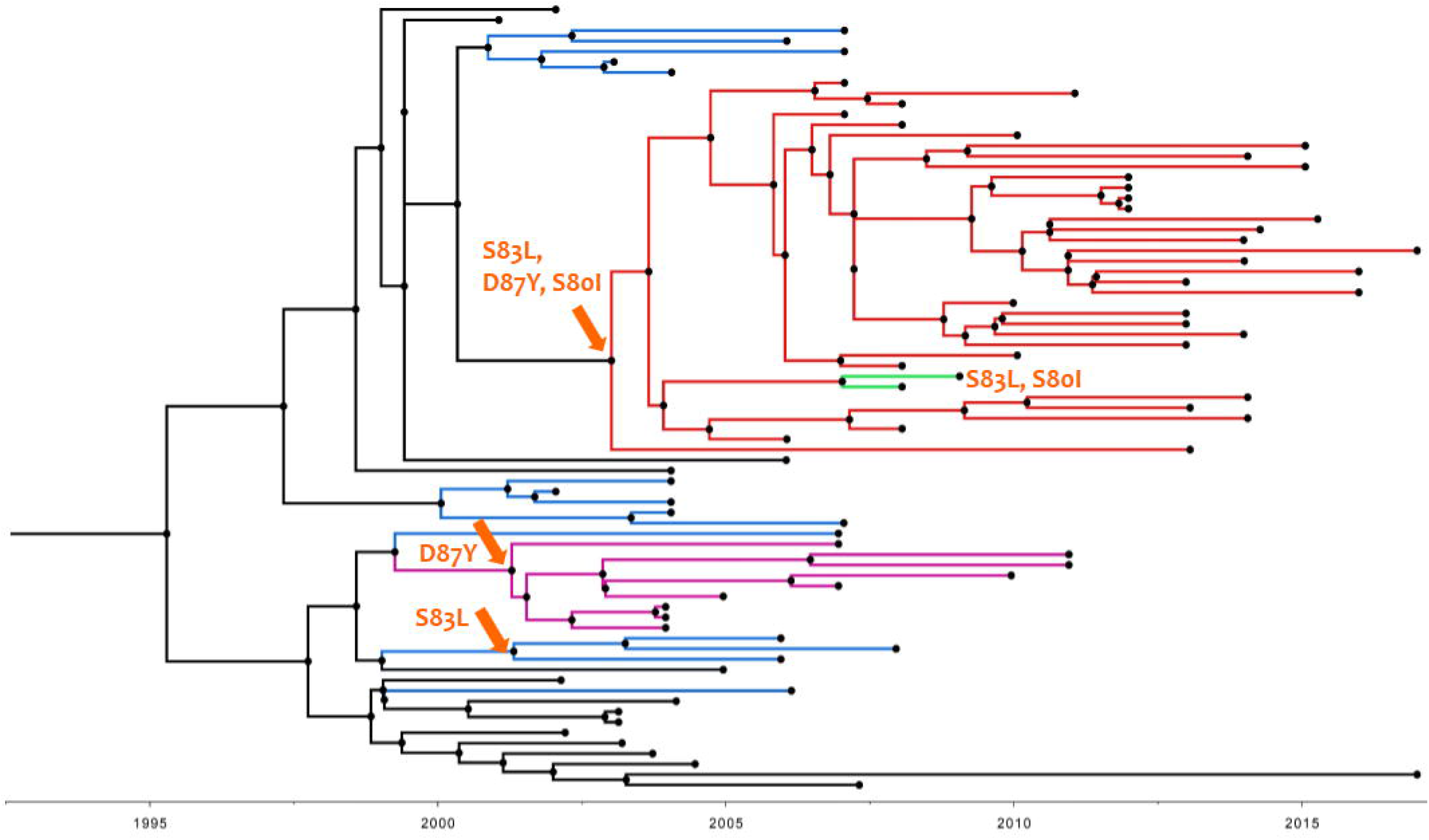
Temporal phylogenetic structure of 97 *S. sonnei* belonging to the Central Asia III lineage from India (1990 – 2017). The orange arrows on the branches indicate the possible occurrence of specific QRDR mutations. Single, double and triple mutations are in different color.

Apart from fluroquinolone resistance, other antimicrobial resistance genes (ARGs) were also identified in Central Asia III lineage. This included genes encoding resistance to previous first-line antimicrobials such as *str*A/B, *tet*A/B, *dfr*A1, *sul*II and aph(3)-lb/aph(6)-id, conferring resistance to streptomycin, tetracyclines, trimethoprim, sulphonamide and aminoglycosides respectively. Other important ARGs were those belonging to the extended-spectrum betalactamases (ESBL) such as *bla*_CTX-M_ and *bla*_TEM_ family. Three *bla*_CTX-M_ and two *bla*_TEM_ variants were identified, this included *bla*_CTX-M-15_, *bla*_CTX-M-14_, *bla*_CTX-M-55_ and; *bla*_TEM-1A_ and *bla*TEM-_1B_ respectively. Also, *mdf*A, an MDR transporter gene was identified. In addition, we found *mph*A gene, a macrolide resistant gene among the Central Asia III *S. sonnei* isolates. Analysis of major plasmids among the isolates showed that Col type, IncB/O/K/Z, IncFIA and IncFIB plasmids were prevalent. Among the Indian Central Asia III *S. sonnei* isolates, genes conferring resistance to first-line antimicrobials and a single MDR transporter gene were widely seen and the Col type plasmid was the most common. The contrasting genomic features and epidemiological distribution of *S. flexneri* and *S. sonnei* are given in **Table 1**.

**Table 1:**
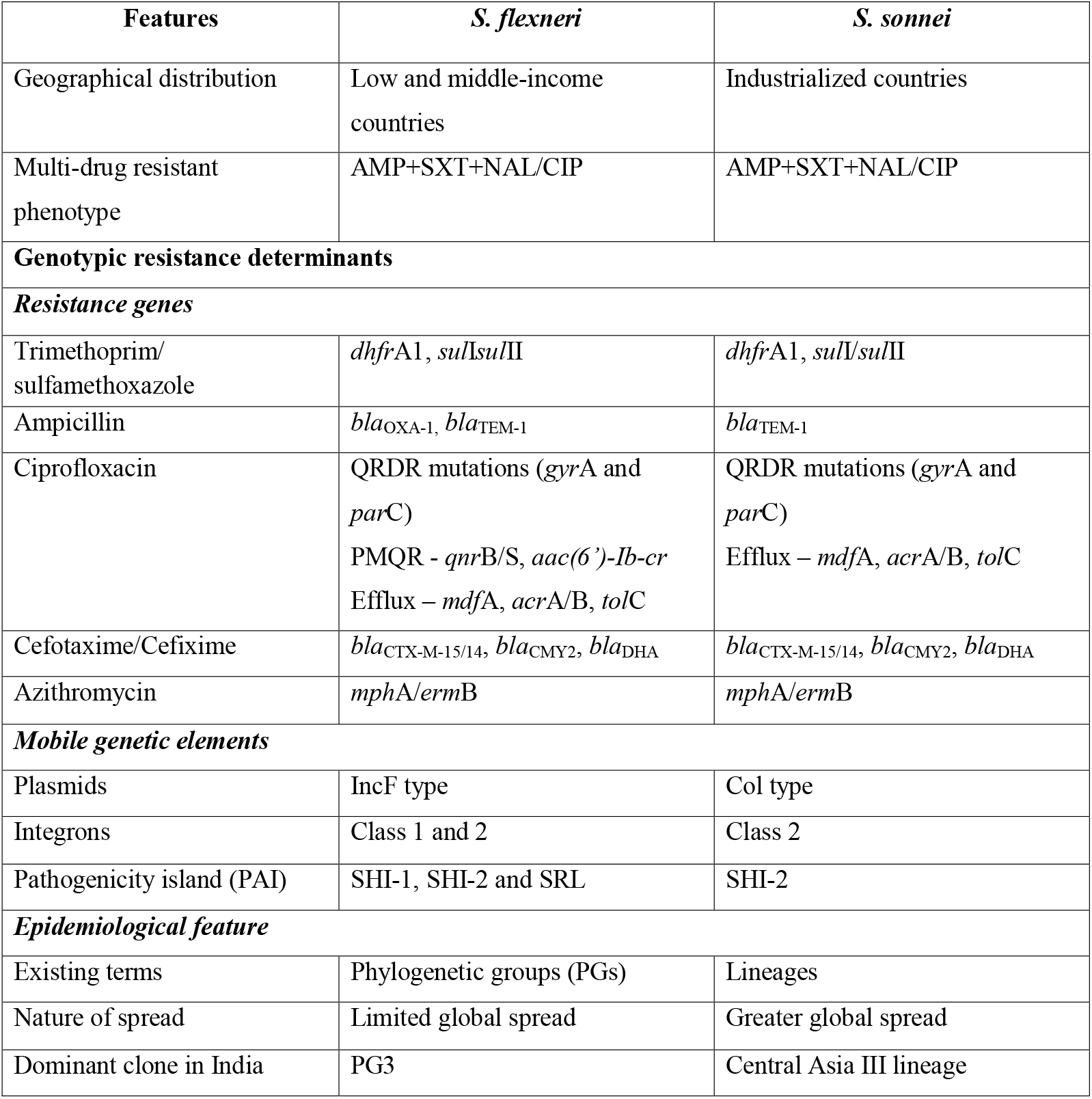
Genomic and epidemiological features of *S. flexneri* versus *S. sonnei*

### Pan-genome analysis

The pan-genome analysis of 106 *S. flexneri* identified 17506 orthologs with 2698 core genes (15%) and 14808 accessory genes (85%). This included, 452 soft core genes, 1628 shell genes and 12728 cloud genes. The total of 9845 unique genes were identified among the *S. flexneri* pan-genome, with FC533 strain belonging to PG3 having the largest number of unique genes (7090). Of these, most of these were found in accessory genome and 26% of them were hypothetical protein.

In *S. sonnei*, the pan-genome analysis for 82 isolates identified 11478 orthologous groups with 662 core genes (6%) and 10816 accessory genes (94%). This included, 689 soft core genes, 3262 shell genes and 6865 cloud genes. Notably, 14 isolates lost a few genes in their core genome and formed a separate cluster in the tree. Majority of these missing genes are found to be hypothetical proteins (54%) with unknown functions and also include various transcriptional regulatory proteins. A total of 4351 unique genes were identified, with highest numbers (2432) identified in FC1793 strain belonging to Central Asia III lineage. Like *S. flexneri*, most of the genes were found in accessory genome and 70% of them were hypothetical protein. The core and accessory gene composition of *S. flexneri* and *S. sonnei* genomes were shown in **(Fig S2 and S3)**.

## Material and Methods

### Bacterial Isolates

106 *S. flexneri* and 82 *S. sonnei* strains isolated from stool specimens from patients with diarrhea or dysentery between the years 1990 and 2017 at the Department of Clinical Microbiology, Christian Medical College, Vellore, India were included in the study. All isolates were retrieved from the archives and confirmed by the standard protocol (Nataro et al., 2011). Serotype identification was done by the slide agglutination test using polyvalent, and monovalent antisera (Denka, Seiken, Japan).

### Antimicrobial susceptibility testing

Antimicrobial susceptibility testing of the isolates against ampicillin (10 μg), trimethoprim/sulphamethoxazole (1.25/23.75 μg), nalidixic acid (30 μg), norfloxacin (10 μg), cefotaxime (30 μg), cefixime (5 μg) and azithromycin (15 μg) was performed using Kirby-Bauer disc diffusion method. The results were interpreted using breakpoints recommended by the Clinical and Laboratory Standards Institute guidelines 2019. The quality control strains used were *E. coli* ATCC 35218 and *E. coli* ATCC 25922.

### Whole genome sequencing

Genomic DNA was extracted from the overnight culture using the QIAmap DNA Mini Kit (Qiagen, Hilden, Germany) as per the manufacturer’s instructions. The quantity and quality of the DNA was analyzed using the Qubit 3.0 fluorometer (Thermofisher, USA) and the Nanodrop spectrophotometer (Thermofisher, USA). Paired-end genomic libraries were prepared with unique indexing of each DNA sample and were sequenced using the short-read Illumina HiSeq V4 platform following the manufacturer’s guidelines.

### SNP based phylogenetic analysis

Phylogenetic analysis of *S. flexneri* and *S. sonnei* genomes were performed against the publically available global genomes. All *S. flexneri* and *S. sonnei* sequences were mapped to the *S. flexneri* 2a str. 301 and the *S. sonnei* Ss046 reference genomes respectively (Accession numbers: NC_007384 and NC_004337.2) using SMALT (version 0.7.4), and SNPs were called against the reference and filtered using SAMtools (Li et al., 2009). This resulted in an alignment of 5, 066 SNPs for *S. sonnei* and 67,454 SNPs for *S. flexneri*, which were used for the phylogenetic inference. A maximum-likelihood (ML) tree was constructed using RAxML v0.7.4 under the GTR substitution model and bootstrap replicates were determined (Stamatakis, 2006).

Temporal signal in the ML phylogeny for *S. sonnei* was investigated using TempEst (http://tree.bio.ed.ac.uk/). The relationship between root to tip distance and the time of isolation were analysed. The temporal phylogenetic structure of *S. sonnei* was determined using the Bayesian Evolutionary Analysis by Sampling Trees, BEAST v.1.10 (Drummond et al., 2012). The recombination free core genome SNP alignment file generated by Gubbins was used as the input to compute the mean evolutionary rate of the genomes and time of the most recent common ancestor (MRCA) (Croucher et al., 2014). Trees were visualized in association with metadata using the web-based interactive Tree of Life (iTOL) (Letunic and Bork, 2019). The study isolates were assigned to previously described lineages or phylogenetic groups (PGs) based on the clustering. The sequences were also analysed for virulence, antimicrobial resistance (AMR), and mobile elements.

### Pan-genome analysis

Pan-genome analysis of *S. flexneri* and *S. sonnei* isolates from India was performed using Roary v. 3.11.2 with default setup (Page et al., 2015). The genomes were annotated using prokka (Seemann, 2014). The genes that were common in all compared strains (core genes) and accessory genes were extracted and was used to contruct a phylogenetic tree. Core genes (99% ≤ strains ≤ 100%), soft core genes (95% ≤ strains < 99%), shell genes (15% ≤ strains < 95%), cloud genes (0% ≤ strains < 15%) and total genes (0% ≤ strains ≤ 100%) were calculated.

### Identification of virulence, resistance and plasmids

Genome data were analyzed for the presence of virulence determinants using VirulenceFinder 1.5 (Joensen et al,. 2014). Acquired antimicrobial resistance genes and chromosomal mutations in the QRDR region were identified using ResFinder 2.1 (Zankari et al., 2012) with a 90% threshold for identity and a 60% of minimum length coverage. The presence of plasmids were analyzed using PlasmidFinder 1.3 (Carattoli et al., 2014) with a 95% threshold for identity.

## Discussion

Here we provide evidence for the long term persistence of all PGs of *S. flexneri* and the recent dominance of ciprofloxacin-resistant *S. sonnei* lineage in India. Unlike *S. sonnei*, replacement of one particular *S. flexneri* PG over the other was not evident, instead we saw that the older PGs have persisted along with the newer ones and continued causing disease as reported earlier. This was evidenced by the fact that every PG comprises at least one isolate collected since early 1990s.

Previously, seven phylogenetic groups (PGs) of *S. flexneri* have been defined by Connor et al., 2015 (Connor et al., 2015). Of the PGs, PG1, 2, 4 and 6 are reported to be the oldest lineages whilst PG3 and PG5 are more recent. In this study, *S. flexneri* isolates were distributed across all the PGs except PG4. The majority of the isolates belonged to PG3 that includes multiple serotypes of *S. flexneri;* however serotype 2 was predominantly seen in this study. This presence of multiple serotypes in a single PG shows that the core genome remains stable but the serotype switching has been common, as reported earlier (Connor et al., 2015). This reassures that unless it is a protein based vaccine carefully calibrated for stability across the phylogeny, vaccines against *Shigella* should continue to target most common serotypes and not to be lineage specific, as happens to be the case.

Our study reveals variability in the composition of virulence factors among *S. flexneri* isolates. Notably, the co-presence of the SHI-1 PAI, which was known to carry three essential virulence genes along with the enterobactin genes and antimicrobial resistance genes were exclusively observed in PG3, which predominantely composed of *S. flexneri* serotype 2a. This may account for the enhanced virulence and international dominance of this serotype. Furhter, AMR has shown to be a strong influence on the recent evolutionary history of many bacterial pathogens (Holt et al., 2012). The sustained presence of resistance genes for first-line antimicrobials in *S. flexneri* over the decades might be an essential factor for the successful maintenance of the lineages in endemic locations including India.

For *S. sonnei*, there are five distinct lineages reported globally. The current pandemic involves globally distributed MDR clones that belong to lineage III. In particular, the ciprofloxacin-resistant Central Asia III lineage are widespread. Although the Central Asia III lineage has spread internationally, its expansion in India has not been previously studied. Our study provides evidence for the dominance of this lineage in India. In addition, a few isolates from Africa, Europe, and America clustered closely with the Indian isolates, which indicates travel related spread. The double/triple mutations in the QRDR in Central Asia lineage III isolates were consistent with previous findings and could be the reason behind increased fitness and global expansion of this lineage alongside the overall synergy between PMQR, *qnr* genes and chromosomal mutations that has been noted in other studies from India (Das et al., 2016). Interestingly, the co-presence of *qnr* genes and mutation in *gyr*A and*par*C genes were identified only in one isolate in our study.

Our data shows temporal introduction of *gyr*A-S83L mutation in 1996 and with the sequential accumulation of secondary mutations leading to the rapid expansion of ciprofloxacin-resistant clones within the Central Asia III lineage in the early 2000s. Our study also indicates that the earliest isolate (1990) within the Central Asia clade was from South Asia, which supports the earlier hypothesis suggesting that South Asia was the likely origin of this lineage (The HC, et al., 2016).

The emergence of resistance to previous first-line antimicrobials such as ampicillin, trimethoprim/sulfamethoxazole and nalidixic acid during this intervening period made ciprofloxacin as the drug of choice to treat drug-resistant shigellosis. Following this, due to the intensive use of ciprofloxacin, resistance to this drug increased from 0.6% in 1998–2000 to 29% in 2007–2009 in Asia and Africa (The HC and Baker, 2018). This further limited the treatment options and made third generation cephalosporins and macrolides as the drugs of choice. With the ESBL producing strains that have left azithromycin as the last-resort (Kotloff et al., 2018, Darton et al., 2018, Salah et al., 2019), the recent emergence of azithromycin resistance in *S. sonnei* and *S. flexneri* serotype 3a, particularly in men who have sex with men (MSM) communities is no surprise (Baker et al., 2015).

Furthermore, the analysis of core and accessory genes of *S. flexneri* and *S. sonnei* revealed variation in composition. *S. sonnei* was found to have larger accessory genome than *S. flexneri*. Strains FC533 and FC1793 was found to have the largest number of unique genes of the *S. flexneri* and *S. sonnei* strains compared respectively. The core genome of *S. flexneri* remains stable suggesting the strength of these genomes in adapting to evolutionary pressures for persistence while, few isolates of *S. sonnei* were missing certain core genes. Earlier studies have shown that the loss of genes is a characteristic of an intracellular pathogenic lifestyle. In *Shigella*, genes that are more prone to deletion are generally associated with pathogenesis and most of this deleted genes are found to be involved in cellular metabolism (Lan and Reeves, 2002). Insights into these genes are out of the scope of this study and can be studied in the future however, it would be interesting to assess whether this gene loss in *S. sonnei* provides any survival benefit to this pathogen.

Reports from previous studies and data from this study show that *Shigella* species from different geographical locations share common AMR and virulence pattern. This indicates that the genetic content of the isolates are merely based on the lineages circulating in the region. The presence of epidemiologically-dominant lineages associated with stable AMR determinants results in the successful survival in the community. The global dissemination of this lineages is more likely facilitated by the frequent travel between other parts of the world.

## Supporting information

The root to tip analysis revealed a strong correlation (R squared 0.7543) between the time of isolation and distance from root suggesting the temporal clock like evolution in the lineage (Fig S1)

The core and accessory gene composition of S. flexneri and S. sonnei genomes were shown in

The core and accessory gene composition of S. flexneri and S. sonnei genomes were shown in

List of genome accession numbers used in this study for phylogenetic analysis

## Supporting information

List of genome accession numbers used in this study for phylogenetic analysis (**Supplementary table**).

## Acknowledgement

We wish to acknowledge the Institutional Review Board of the Christian Medical College, Vellore (83-i/11/13) for approving the study. We thank the sequencing team at the Wellcome Trust Sanger Institute for sequencing the samples. The study was supported by the Indian Council of Medical Research, New Delhi (Ref. No: AMR/TF/55/13ECDII dated 23/10/2013).

## Supplementary Figure Legends

**Fig S1:** Root-to-tip branch lengths extracted from the maximum-likelihood tree of *S. sonnei* are plotted against the year of isolation

**Fig S2:** The core and accessory gene composition of 106 *S. flexneri* isolates extracted using Roary. 2698 core genes were shared by all strains while unique genes identified were 9845.

**Fig S3:** The core and accessory gene composition of 82 *S. sonnei* isolates extracted using Roary.662 core genes were shared by all strains while 4351unique genes were identified.

