## Supplementary figures and images for "Phylogenetic and evolutionary analysis reveals the recent dominance of ciprofloxacin resistant *S. sonnei* and local persistence of *S. flexneri* clones in India"

### The core and accessory gene composition of S. flexneri and S. sonnei genomes were shown in

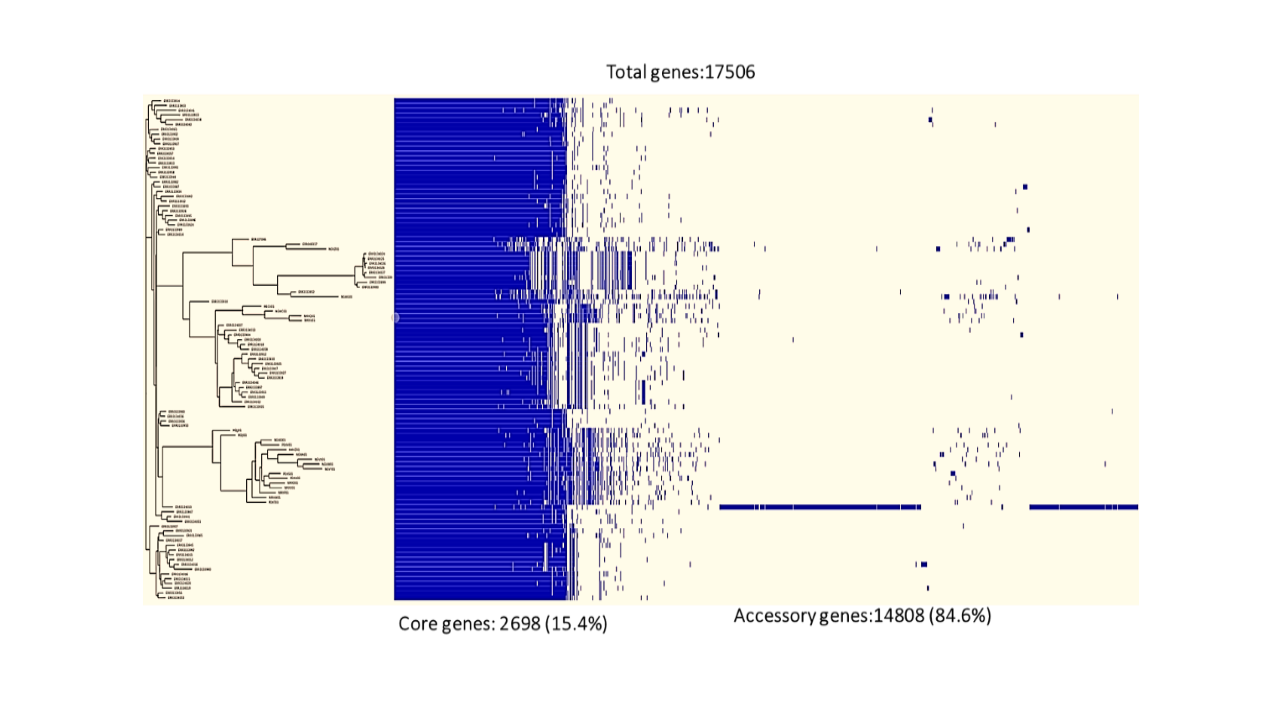

### The core and accessory gene composition of S. flexneri and S. sonnei genomes were shown in

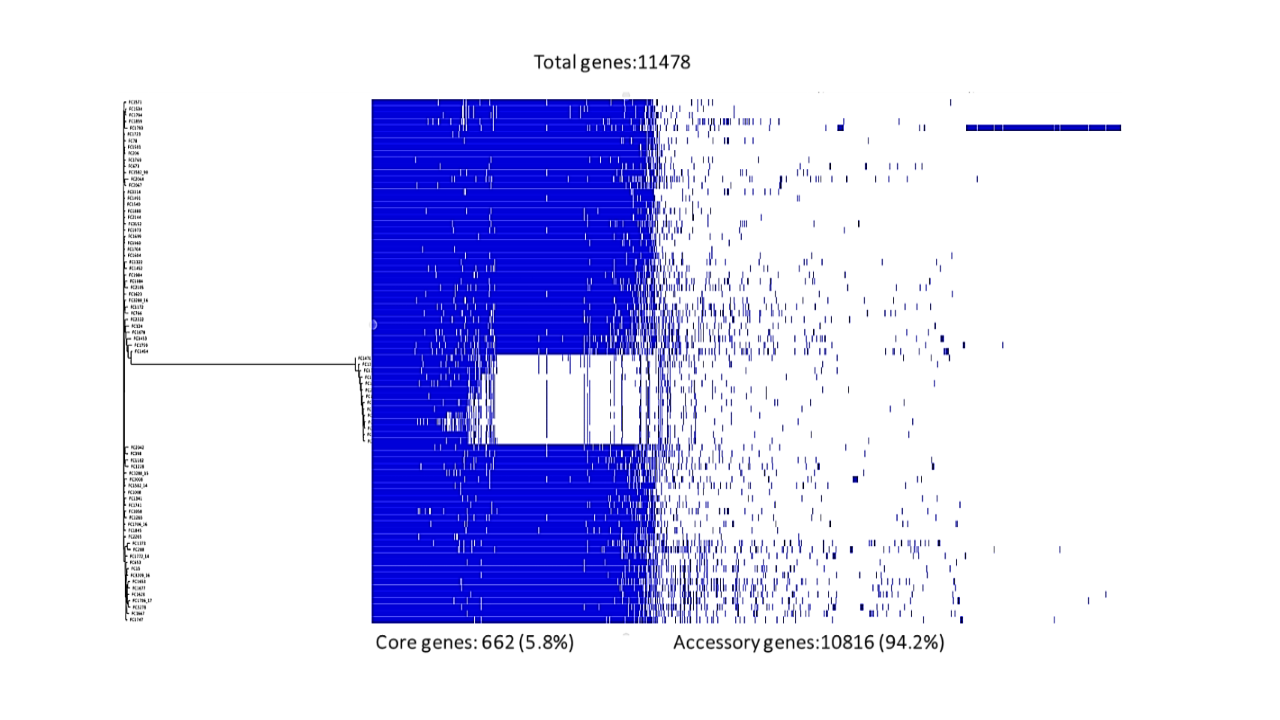

### The root to tip analysis revealed a strong correlation (R squared 0.7543) between the time of isolation and distance from root suggesting the temporal clock like evolution in the lineage (Fig S1)

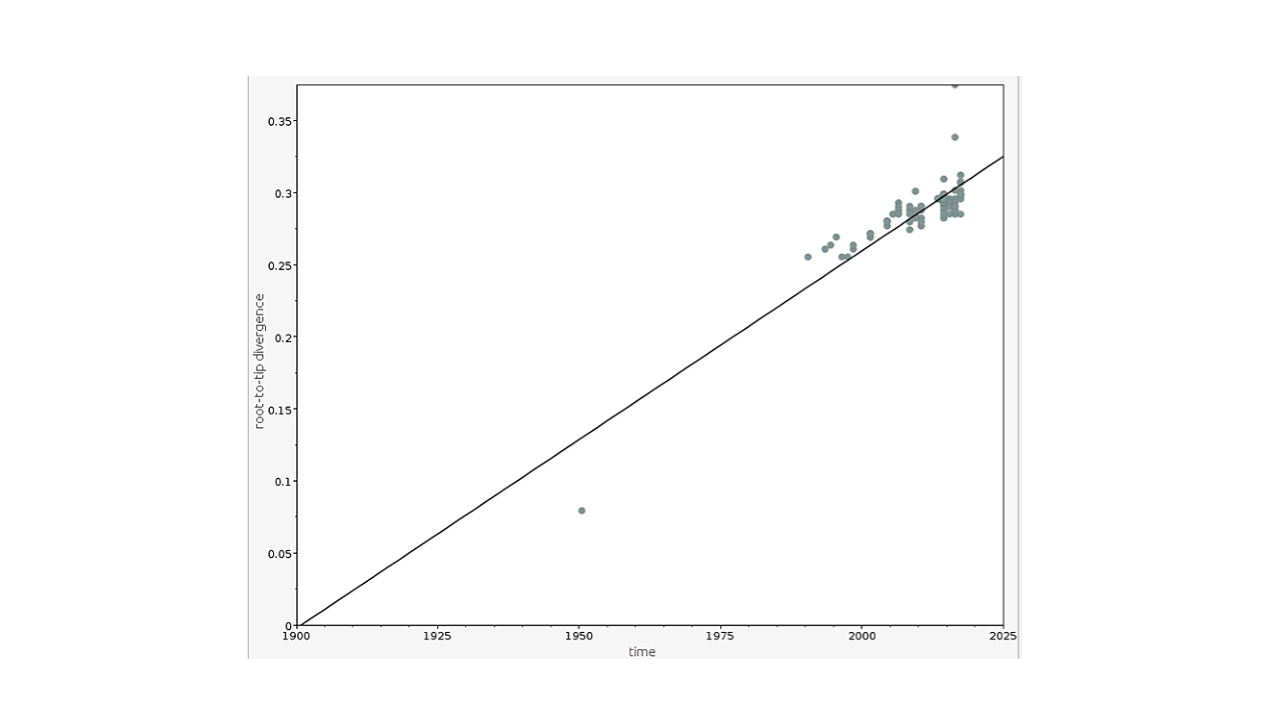
