## Supplementary material for "Phylogenetic and evolutionary analysis reveals the recent dominance of ciprofloxacin resistant *S. sonnei* and local persistence of *S. flexneri* clones in India": List of genome accession numbers used in this study for phylogenetic analysis

**Table S1:** List of *Shigella sonnei* isolates sequenced and analyzed in this study

| **Sample ID** | **Accession No.** |  | **Sample ID** | **Accession No.** |  | **Sample ID** | **Accession No.** |
| --- | --- | --- | --- | --- | --- | --- | --- |
| FC288 | NGWI00000000 |  | FC1960 | ERR3133972 |  | FC1228 | ERR3134052 |
| FC1373 | NGWH00000000 |  | FC1845 | ERR3133973 |  | FC2105 | ERR3134054 |
| FC1772 | NGWB00000000 |  | FC2265 | ERR3133974 |  | FC1984 | ERR3134055 |
| FC3278 | NMYB00000000 |  | FC1880 | ERR3133975 |  | FC2222 | ERR3134056 |
| FC653 | NMXY00000000 |  | FC1534 | ERR3133976 |  | FC1974 | ERR3134058 |
| FC3209 | NMXU00000000 |  | FC3209 | ERR3133978 |  | FC1172 | ERR3134060 |
| FC1747 | NMXS00000000 |  | FC1470 | ERR3133979 |  | FC1772 | ERR3133977 |
| FC15 | NMXR00000000 |  | FC2127 | ERR3133980 |  | FC1582 | ERR3133969 |
| FC1706 | PDYB00000000 |  | FC1454 | ERR3133995 |  | FC1677 | ERR3133959 |
| FC1628 | PDYA00000000 |  | FC1759 | ERR3133996 |  |  |  |
| FC1667 | PDXZ00000000 |  | FC3288 | ERR3133997 |  |  |  |
| FC1653 | PDXX00000000 |  | FC1772 | ERR3133998 |  |  |  |
| FC1677 | PDXW00000000 |  | FC1265 | ERR3133999 |  |  |  |
| FC1582 | ERR3133891 |  | FC1706 | ERR3134001 |  |  |  |
| FC1769 | ERR3133904 |  | FC3314 | ERR3134002 |  |  |  |
| FC1723 | ERR3133906 |  | FC3152 | ERR3134003 |  |  |  |
| FC1571 | ERR3133918 |  | FC3008 | ERR3134004 |  |  |  |
| FC673 | ERR3133920 |  | FC3306 | ERR3134005 |  |  |  |
| FC78 | ERR3133922 |  | FC3288 | ERR3134008 |  |  |  |
| FC206 | ERR3133928 |  | FC2973 | ERR3134009 |  |  |  |
| FC1501 | ERR3133929 |  | FC766 | ERR3134021 |  |  |  |
| FC1704 | ERR3133937 |  | FC1231 | ERR3134022 |  |  |  |
| FC1491 | ERR3133938 |  | FC1084 | ERR3134023 |  |  |  |
| FC1452 | ERR3133939 |  | FC2042 | ERR3134029 |  |  |  |
| FC1741 | ERR3133940 |  | FC2064 | ERR3134030 |  |  |  |
| FC1008 | ERR3133941 |  | FC2067 | ERR3134031 |  |  |  |
| FC1058 | ERR3133943 |  | FC920 | ERR3134033 |  |  |  |
| FC1859 | ERR3133960 |  | FC956 | ERR3134034 |  |  |  |
| FC1540 | ERR3133961 |  | FC1453 | ERR3134035 |  |  |  |
| FC1793 | ERR3133962 |  | FC398 | ERR3134039 |  |  |  |
| FC1794 | ERR3133963 |  | FC1182 | ERR3134040 |  |  |  |
| FC2144 | ERR3133964 |  | FC1078 | ERR3134043 |  |  |  |
| FC1582 | ERR3133965 |  | FC124 | ERR3134044 |  |  |  |
| FC1623 | ERR3133966 |  | FC1316 | ERR3134045 |  |  |  |
| FC1604 | ERR3133967 |  | FC1219 | ERR3134047 |  |  |  |
| FC1699 | ERR3133968 |  | FC1322 | ERR3134048 |  |  |  |
| FC1975 | ERR3133970 |  | FC1341 | ERR3134049 |  |  |  |
| FC1973 | ERR3133971 |  | FC1353 | ERR3134050 |  |  |  |

**Table S2:** List of *Shigella flexneri* isolates sequenced and analyzed in this study

| **Sample ID** | **Accession No.** |  | **Sample ID** | **Accession No.** |  | **Sample ID** | **Accession No.** |
| --- | --- | --- | --- | --- | --- | --- | --- |
| FC1180 | MDJJ00000000 |  | FC1966 | ERR3134016 |  | FC1050 | ERR3133926 |
| FC1139 | MECX00000000 |  | FC3257 | ERR3134017 |  | FC909 | ERR3133903 |
| FC1172 | MDJI00000000 |  | FC3090 | ERR3134018 |  | FC1251 | ERR3133909 |
| FC1417 | NGWG00000000 |  | FC566 | ERR3134019 |  | FC1739 | ERR3133902 |
| FC906 | NGWD00000000 |  | FC929 | ERR3134020 |  | FC122 | ERR3133932 |
| FC1182 | NGWC00000000 |  | FC1885 | ERR3134024 |  | FC409 | ERR3133917 |
| FC1659 | NGWA00000000 |  | FC2013 | ERR3134025 |  | FC3513 | ERR3133985 |
| FC470 | NGVZ00000000 |  | FC1996 | ERR3134026 |  | FC3421 | ERR3133989 |
| FC1247 | NGVY00000000 |  | FC1800 | ERR3134027 |  | FC6_2010 | ERR3133935 |
| FC1607 | NGVX00000000 |  | FC1783 | ERR3134028 |  | FC1255 | ERR3133947 |
| FC1481 | NGVW00000000 |  | FC425 | ERR3134032 |  | FC3056 | ERR3133956 |
| FC3433 | NMXZ00000000 |  | FC1673 | ERR3134036 |  | FC1268 | ERR3133954 |
| FC1170 | NMXX00000000 |  | FC1282 | ERR3134038 |  | FC1049 | ERR3133953 |
| FC1824 | NMXW00000000 |  | FC26 | ERR3134041 |  | FC13_rpt | ERR3133893 |
| FC601 | NMXV00000000 |  | FC1997 | ERR3134042 |  | FC1506 | ERR3133950 |
| FC401 | NMXQ00000000 |  | FC1788 | ERR3134046 |  | FC3017 | ERR3133957 |
| FC420 | NMXP00000000 |  | FC1310 | ERR3134051 |  | FC3442 | ERR3133983 |
| FC248 | NMXO00000000 |  | FC2145 | ERR3134053 |  | FC2007 | ERR3133933 |
| FC1405 | PDXV00000000 |  | FC2003 | ERR3134057 |  | FC2094 | ERR3133987 |
| FC2101 | PDXU00000000 |  | FC2188 | ERR3134059 |  | FC3063 | ERR3133944 |
| FC2414 | PDXT00000000 |  | FC1604 | ERR3134061 |  | FC108 | ERR3133936 |
| FC1954 | PDXS00000000 |  | FC2947 | ERR3133952 |  | FC1461 | ERR3133990 |
| FC393 | ERR3133892 |  | FC2463 | ERR3133948 |  | FC1488 | ERR3133982 |
| FC13 | ERR3133893 |  | FC6_1999 | ERR3133931 |  | FC2917 | ERR3133958 |
| FC1728 | ERR3133895 |  | FC2595 | ERR3133919 |  | FC1570 | ERR3133992 |
| FC1876 | ERR3133896 |  | FC1113 | ERR3133927 |  | FC2927 | ERR3133991 |
| FC315 | ERR3133897 |  | FC2216 | ERR3133925 |  | FC3000 | ERR3133945 |
| FC1905 | ERR3133899 |  | FC848 | ERR3133907 |  |  |  |
| FC1793 | ERR3133900 |  | FC1834 | ERR3133912 |  |  |  |
| FC1763 | ERR3134000 |  | FC637 | ERR3133949 |  |  |  |
| FC2899 | ERR3134006 |  | FC1536 | ERR3133911 |  |  |  |
| FC3085 | ERR3134007 |  | FC2004 | ERR3133984 |  |  |  |
| FC533 | ERR3134010 |  | FC1694 | ERR3133930 |  |  |  |
| FC3124 | ERR3134011 |  | FC1277 | ERR3133910 |  |  |  |
| FC1912 | ERR3134012 |  | FC2245 | ERR3133921 |  |  |  |
| FC1550 | ERR3134013 |  | FC2242 | ERR3133923 |  |  |  |
| FC563 | ERR3134014 |  | FC193 | ERR3133924 |  |  |  |
| FC3056_16 | ERR3134015 |  | FC401_95 | ERR3133914 |  |  |  |
